# Adaptive and maladaptive consequences of deregulation in a bacterial gene regulatory network

**DOI:** 10.1101/2023.01.30.526227

**Authors:** Rhea Vinchhi, Chetna Yelpure, Manasvi Balachandran, Nishad Matange

## Abstract

The archetypal PhoQP two-component system from Enterobacteria regulates crucial pathways like magnesium homeostasis in *Escherichia coli* and virulence factor expression in *Salmonella enterica*. Previously we had reported that a laboratory strain of *E. coli* rapidly accumulated loss-of-function mutations in the *mgrB* gene, a negative feedback regulator of PhoQP, when evolved in the presence of the antibiotic trimethoprim. Hyperactive PhoQP enhanced the expression of dihydrofolate reductase (*folA*), target of trimethoprim, resulting in antibiotic tolerance. Here we ask, firstly, how important are mutations in *mgrB* for trimethoprim resistance? Using laboratory evolution, we show that trimethoprim resistance evolves by different mutational trajectories under condition of high and low PhoQP activity. Mutations in *mgrB* are only fixed when PhoQP is active. Importantly, loss of functional MgrB, though itself only mildly beneficial, enhances the fixation probability of trimethoprim-resistant bacteria under selection and this can be explained by epistasis between *mgrB* and *folA* loci. As a result, the activation status of PhoQP directly impacts how fast resistance is acquired by evolving populations of *E. coli*. Secondly, we investigate why negative feedback may be needed in the PhoQP system. We show that under drug-free conditions MgrB is required to mitigate the fitness costs of pervasive gene dysregulation by hyperactive PhoQP. Using RNA-seq transcriptomics and genetic analyses, we demonstrate that PhoQP-hyperactivation perturbs the balance of RpoS and RpoD-regulated transcriptional programs, and spontaneous mutations in *rpoS* rectify this imbalance. We propose that deregulation can be adaptive or maladaptive depending on the environmental context and this explain the evolution of negative feedback in bacterial gene regulatory networks.

## Introduction

Signal transduction pathways perform the vital function of altering cellular physiology in response to the environment. Signaling cascades often culminate in gene expression states that dictate the behavior of cells and facilitate adaptation to stressful environments. Biochemical and genetic experiments have elucidated the roles of individual signaling cascades in detecting stimuli at the cell surface, transducing this signal intracellularly and modulating gene expression by changing activation states of transcription factor/s. Building on this information, systems-level approaches have revealed that signaling proteins are organized into networks, often referred to as gene regulatory networks [1, 2]. Organization of regulatory proteins into networks not only brings in robustness and exquisite control, but also allows for crosstalk between signaling cascades, ultimately shaping which genes are expressed and to what extent [1-3].

A ubiquitous feature of gene regulatory networks, seen from bacteria to humans, is the presence of feedback [1, 2, 4-7]. Positive feedback in regulatory networks ensures rapid ‘switch-like’ behavior in response to an activating signal [7, 8]. It allows for bistability, i.e., the existence of two relatively stable transcriptional states [9]. In Gram positive bacteria such as *Bacillus subtilis*, bistability plays an important role in governing cell cycle and developmental programs like sporulation [9]. Negative feedback on the other hand dampens, restricts, or resets activation levels of a signaling network. Negative feedback is thought to be corrective and is required to calibrate levels of gene expression and buffer noise [3, 10, 11]. Corrective feedback has been shown to facilitate adaptation in bacterial signaling as well as linearize gene expression responses [12].

The major signaling proteins used by bacteria are two-component systems [13-15]. The first component, referred to as ‘sensor kinase’, auto-phosphorylates at a conserved histidine residue upon activation. It then transfers its phosphate group to a conserved aspartate residue on a second protein, called the ‘response regulator’, which is usually a transcription factor itself. Phosphorylation status of the response regulator directly alters its ability to bind to gene promoters and modify gene expression [13, 15]. Several variations on this basic schema are observed in nature, including dual kinase-phosphatase activity of the sensor kinase [16], multiple phosphorelay steps [13, 17] and crosstalk between non-cognate kinases and response regulators [18, 19]. Two-component systems across bacteria orchestrate responses to a wide range of cues such as metal ions [20], pH [21], nutrients [22], antimicrobial peptides [23], quorum sensing molecules [24] and cell envelope stress [25, 26]. More recently, two-component sensor kinases have been proposed as novel targets for the design of antimicrobials owing to their association with drug resistance in several clinically relevant bacterial species [27-29].

Two-component signaling pathways are important substrates for adaptive evolution in bacteria and mutations in two-component signaling proteins are responsible for adaptation to adverse environments. For instance, sequence variation in the EvgAS two-component system is implicated in modulating the survival of *Escherichia coli* strains in low pH [30]. Similarly, mutations in the DosR-DosS-DosT system in the Beijing lineage of *Mycobacterium tuberculosis* have been associated with hypervirulence [31, 32]. We showed previously that loss-of-function mutations in the *mgrB* gene are rapidly enriched in *E. coli* bacteria that were exposed to the antibiotic trimethoprim [33]. The *mgrB* gene codes for a small membrane protein that inhibits the PhoQP two-component system by binding to the sensor kinase PhoQ [34, 35]. Expression of *mgrB* is itself activated by PhoP, setting up a negative feedback loop [34, 36, 37]. Loss of functional MgrB resulted in PhoQP-dependent overproduction of dihydrofolate reductase (DHFR) [33, 38], which conferred tolerance to trimethoprim [33]. Interestingly, loss of functional *mgrB* in clinical strains of *Klebsiella pneumoniae* [39] and *Enterobacter cloacae* [40] is causally linked to resistance to colistin, a last-resort antibiotic, demonstrating that deregulation of PhoQP can be beneficial in different bacterial species under antibiotic pressure. In this study, we build on our earlier observations to investigate how crucial mutational loss of negative feedback in PhoQP is for the evolution of trimethoprim resistance in *E. coli*, and why negative feedback may have evolved and is maintained by two-component signaling pathways.

## Results

### Mutations in *mgrB, folA* and *rpoS* genes drive adaptation to trimethoprim in laboratory-evolved *E. coli*

In our laboratory evolution experiments loss of functional *mgrB* was the first mutational event in *E. coli* adapting to trimethoprim [33]. Long term antibiotic exposure led to adaptive sweeps involving mutations in 2 other genes i.e., *folA*, which codes for dihydrofolate reductase (DHFR) and *rpoS*, which codes for the enterobacterial stationary phase sigma factor [33]. While mutations in *mgrB* and *folA* directly enhanced trimethoprim resistance, *rpoS*-mutations enhanced fitness without altering drug IC_50_ [33]. To test whether these 3 loci reproducibly accumulated mutations during the evolution of trimethoprim resistance, we sequenced the genomes of 3 randomly picked resistant isolates from 3 independent lineages of *E. coli* after 350 generations of evolution in trimethoprim. Mutations in *folA* were found in isolates from 2 of the 3 lineages, while *mgrB* harboured mutations in all 3 lineages (Figure 1, Supplementary File 1). Like *mgrB, rpoS* mutations too were found across all 3 lineages (Figure 1, Supplementary File 1). Though mutations in several other genes were also found (Supplementary File 1), *folA, mgrB* and *rpoS* were the only loci that harboured mutations consistently across lineages, reinforcing their role as the primary hotspots for adaption to trimethoprim.

**Figure 1.**
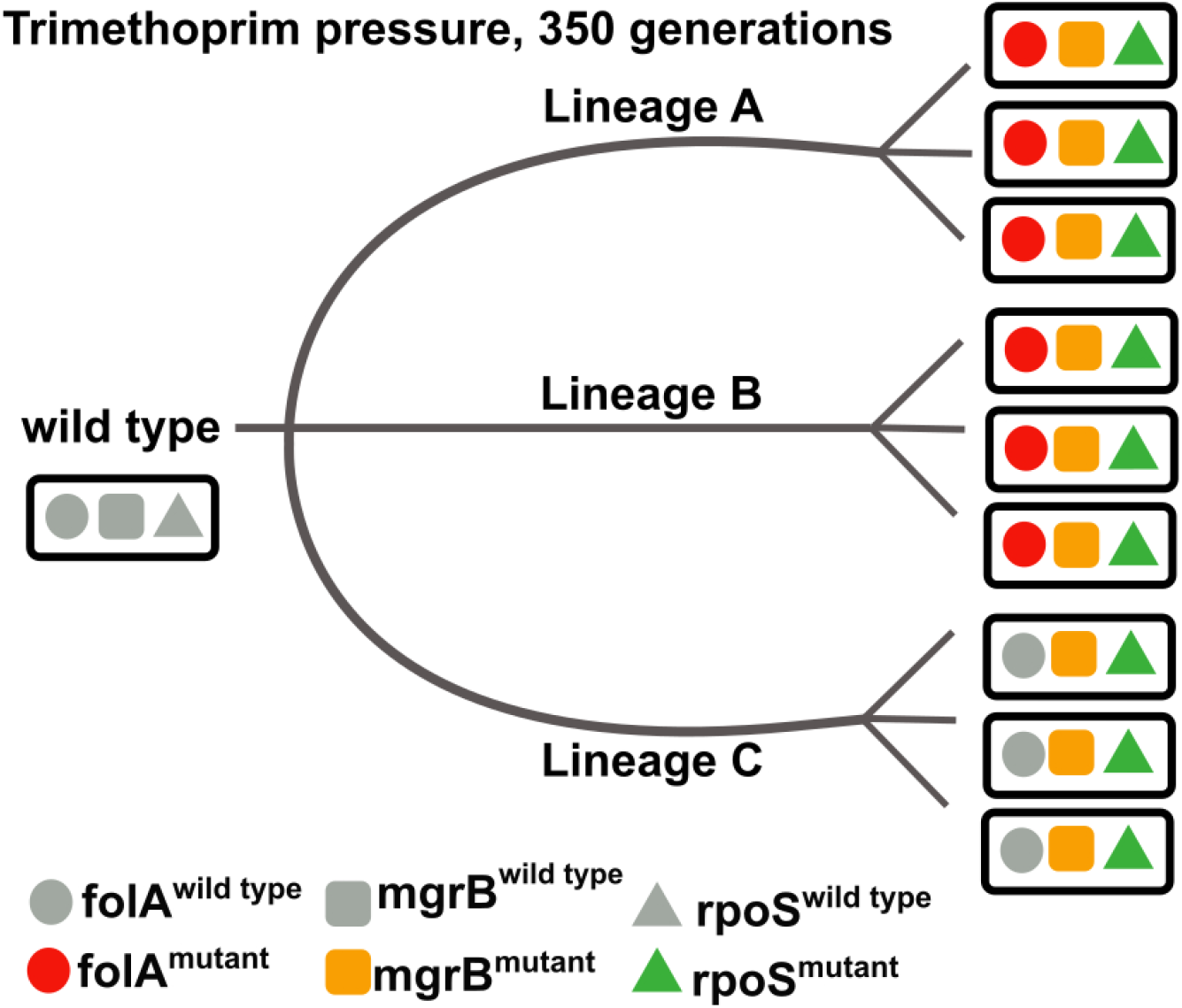
Genetic changes associated with trimethoprim-resistance evolution. Diagrammatic representation of three gene loci, *folA* (circle), *mgrB* (square) and *rpoS* (triangle) across 3 randomly selected trimethoprim-resistant isolates from 3 independently evolved lineages (A, B and C) [33]. Each lineage was subjected to sub-MIC trimethoprim (300 ng/mL) for ∼350 generations after which resistant bacteria were isolated on media supplemented with 1 µg/mL of trimethoprim (i.e. MIC of wild type). Wild type *folA, mgrB* and *rpoS* are represented in gray, while coloured symbols are used to represent mutant loci. All mutations in *mgrB* and *rpoS* are known/predicted to be loss of function mutations. Mutations in *folA* were in the coding region or the gene promoter. A complete list of mutations detected in each of the isolates is given in Supplementary File 1.

### Mutational loss of *mgrB* facilitates evolution of trimethoprim resistance

By itself, loss of *mgrB* led to a mild enhancement in drug IC_50_ [33]. Despite this, *mgrB* was consistently mutated across independently evolved trimethoprim-resistant bacteria (Figure 1). This observation suggested that though *mgrB*-deficiency alone had only a small effect on drug resistance, it may serve a facilitatory role during adaptation to trimethoprim. To empirically test this idea, we asked, firstly, whether it was possible for *E. coli* to evolve resistance without implicating *mgrB* and secondly, if resistance did evolve without mutations in *mgrB*, how it would impact the rate of resistance evolution.

In strain competition experiments, *E. coli ΔmgrB* showed a dose-dependent increase in relative fitness in trimethoprim-supplemented media (Figure 2A). High concentration of Mg^2+^ (5-10 mM) in growth media, which is known to inhibit PhoQ activity [21, 41] completely neutralized this fitness advantage of *mgrB*-deficiency (Figure 2A). Thus, loss of MgrB was only advantageous for *E. coli* under conditions in which PhoQP was active. Based on this result, we established 6 evolving lineages (1, 2, 3: High Mg^2+^; 4, 5, 6: Low Mg^2+^) that were exposed to sub-MIC trimethoprim (100 ng/mL) under conditions of high or low PhoQP activity for ∼210 generations (Figure 2B). We argued that, since PhoQP would be active only in low Mg^2+^, these lineages alone would evolve mutations in *mgrB*, allowing us to dissect out its role in the evolution of resistance. In all three Low Mg^2+^ lineages, trimethoprim-resistant bacteria were rapidly fixed (i.e. frequency >0.9) within the first 50 generations of evolution (Figure 2C). In contrast, High Mg^2+^ lineages evolved resistance at much slower rates, and resistant bacteria were unable to reach fixation over the duration of the experiment (Figure 2C). To ensure that this effect was due to the activity of PhoQP and not an independent effect of Mg^2+^, we established 3 independent evolving lineages starting with an isogenic *E. coli ΔphoP* strain in low Mg^2+^ media supplemented with trimethoprim (Figure 2B). In these populations too, trimethoprim-resistant bacteria did not get fixed and remained at low frequencies even after 200 generations of evolution (Figure 2C). Thus, low/no active PhoQP severely impeded the establishment of trimethoprim-resistance in *E. coli* populations.

**Figure 2.**
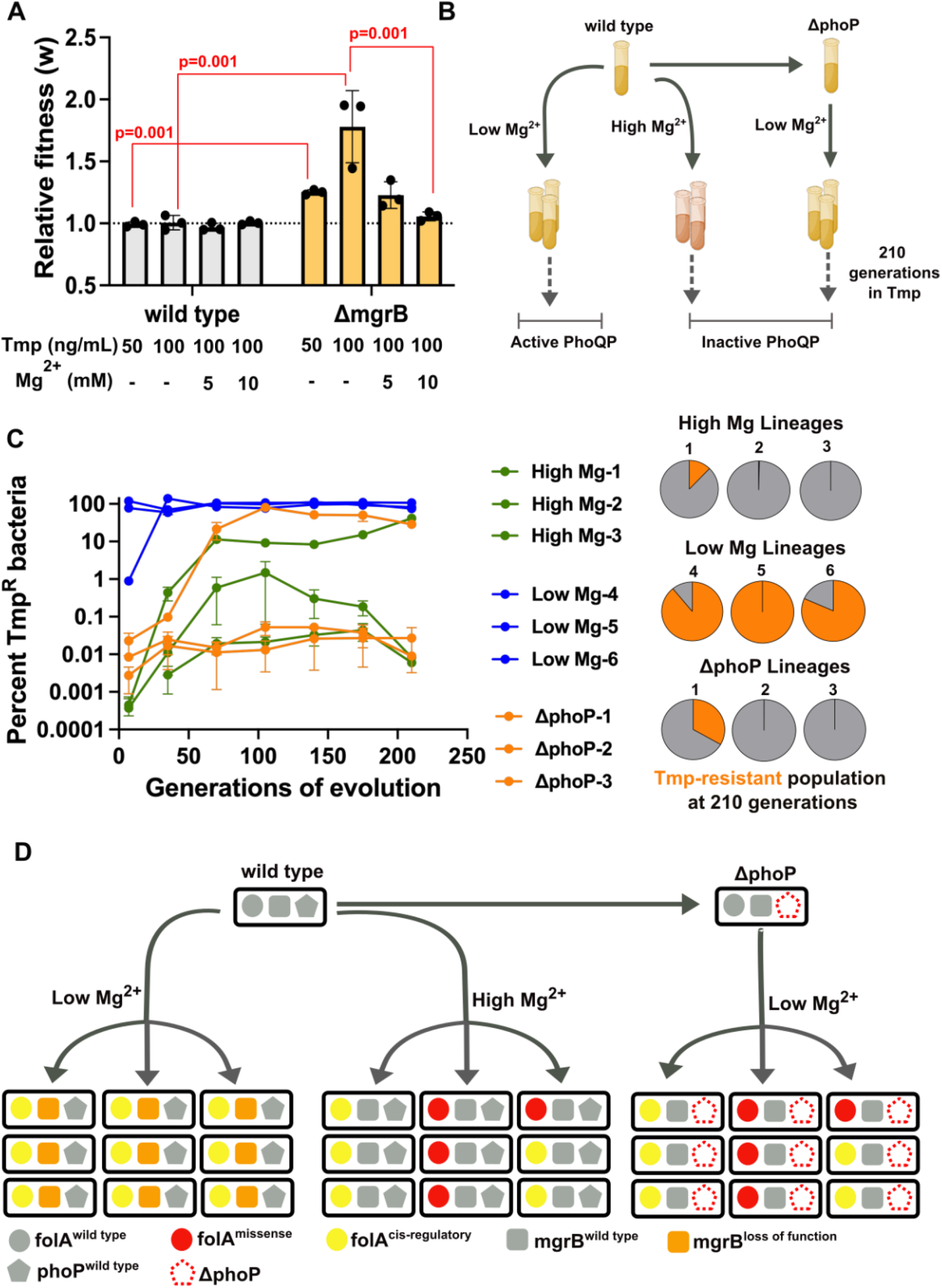
Loss of MgrB facilitates the evolution of trimethoprim resistance in *E. coli*. **A**. Relative fitness (w) of *E. coli* wild type (gray) and an *mgrB*-knockout (orange) strain compared to an isogenic *lacZ*-knock out reference strain. Relative fitness was calculated in different concentrations of trimethoprim (Tmp) and MgSO_4_ as indicated. Individual replicates are plotted as black circles, and the mean value is shown as a bar. Error bars represent standard deviation. A value of 1 (dotted line) indicates no change in fitness compared to the reference strain. Statistical significance was tested using a Student’s t-test and p-values are shown for relevant comparisons. **B**. Schematic for laboratory evolution to investigate the contribution of *mgrB*-mutations to trimethoprim resistance. Triplicate lineages were evolved for ∼210 generations in media supplemented with sub-MIC trimethoprim (100 ng/mL) in the presence or absence of 10 mM MgSO_4_. An additional 3 lineages were also evolved starting with an isogenic *ΔphoP* strain in the absence of MgSO_4_. The status of PhoQP activity in each of the conditions is indicated. **C**. Numbers of trimethoprim-resistant bacteria from evolving populations shown in B. Left panel: Titers of resistant bacteria at different generations of evolution under active and inactive PhoP conditions expressed as a percentage of total population. Right panel: Resistant fraction for each of the 9 lineages at 210 generations of evolution (orange) shown as pie charts. **D**. Diagrammatic summary of genome sequencing for 3 randomly picked trimethoprim-resistant isolates from each of the 9 lineages in B and C. The *folA* gene is represented by a circle, *mgrB* by a square and *phoP* by a pentagon. Gray filled symbols indicate wild type alleles, while coloured symbols indicate mutations as shown in the key.

Genome re-sequencing of 3 randomly picked resistant isolates from each of the lineages confirmed that, as expected, mutations in *mgrB* occurred in all Low Mg^2+^ isolates (Figure 2D, Supplementary File 2). In contrast, none of the isolates from High Mg^2+^ and *ΔphoP* lineages harboured *mgrB*-mutations (Figure 2D, Supplementary File 2). Importantly, all 27 sequenced trimethoprim-resistant isolates harboured mutations at the *folA* locus, either in the gene promoter or in the coding sequence (Figure 2D, Supplementary File 2). Genome sequencing of the entire population further confirmed these findings (Supplementary File 2). These results demonstrated that mutations in *folA* were sufficient for the evolution of trimethoprim-resistance. However, *mgrB*-mutations, though not necessary for the evolution of resistance, facilitated fixation of resistant bacteria under trimethoprim selection.

### Epistasis between *mgrB* and *folA* facilitates the fixation of trimethoprim resistant bacteria

We next investigated the mechanism underlying the facilitatory role played by *mgrB*-loss. Intergenic epistasis is a well-known determinant of the mutational landscape of resistant bacteria evolving at sub-MIC drug [42]. Therefore, we first assessed whether mutations in *mgrB* were epistatic with drug-resistant *folA* alleles. For these analyses we chose representative strains that harboured mutations in *mgrB* alone, *folA* alone (coding region and its promoter) or at both loci and calculated fold-IC_50_ values over wild type. Loss of *mgrB* alone (*E. coli ΔmgrB*) resulted in a marginal enhancement in IC_50_ of ∼3-fold over wild type (Figure 3A). On the other hand, isolates harbouring either a missense mutation in the *folA* gene (Isolate Tmp^R^-*folA*-Trp_30_Arg) or a promoter-up mutation (Isolate Tmp^R^-*folA*-C_-35_T) had significantly higher values of fold-IC_50_ (Figure 3A). Based on a non-epistatic/additive model [42], we expected ∼152-fold or ∼41-fold increase in IC_50_ of strains harbouring *mgrB*-mutations in addition to the *folA*-C_-35_T and *folA*-Trp_30_Arg alleles respectively (Figure 3A). Deviation from these expected values would indicate epistasis between *mgrB* and *folA*. We selected resistant isolates Tmp^R^-A and Tmp^R^-B, derived from our long-term evolution lines to test this prediction. Tmp^R^-A harboured mutant *mgrB* and *folA*-C_-35_T, while Tmp^R^-B harboured mutant *mgrB* and *folA*-Trp_30_Arg. Interestingly, IC_50_ values of both isolates deviated from expectation, but in opposite ways. Though both strains had higher IC_50_ values than *mgrB*/*folA* mutants alone, Tmp^R^-A showed “less-than-additive” effect or magnitude epistasis, while Tmp^R^-B showed “greater-than-additive” or synergistic epistasis (Figure 3A). To verify that the observed effects could be attributed to hyperactive PhoQP we deleted the *phoP* gene from Tmp^R^-A and Tmp^R^-B. Indeed, deleting *phoP* from Tmp^R^-A led to only a ∼10 % reduction in fold-IC_50_ while Tmp^R^-B showed close to ∼80 % reduction in fold-IC_50_ (Figure 3A), corroborating our results. These experiments showed that mutations in *mgrB* enhanced the IC_50_ values of bacteria harbouring *folA* mutations, but this effect was more pronounced for coding mutations than promoter mutations.

**Figure 3.**
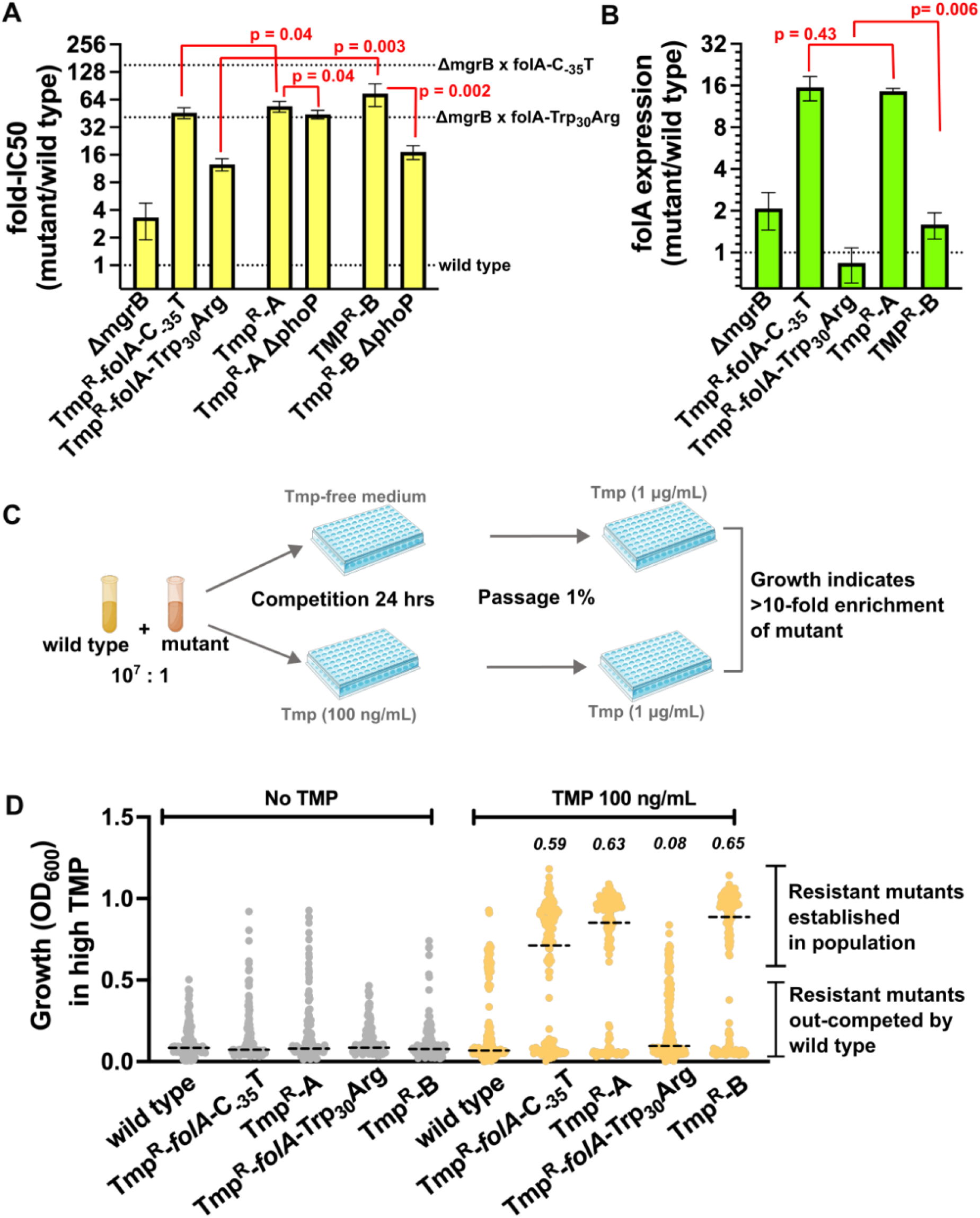
Epistasis between *mgrB* and *folA* mutations facilitate fixation of resistant bacteria. **A**. Fold change in trimethoprim IC_50_ values over wild type *E. coli* for indicated resistant strains are plotted as mean ± SD from 3 independent measurements. Statistical significance was tested using a Student’s t-test and p-values for relevant comparisons are shown. Expected values for indicated combinations of mutations based on a non-epistatic/additive model are shown as dotted lines. **B**. Expression level of the *folA* gene measured by quantitative RT-PCR in trimethoprim resistant strains, normalized to wild type (=1; dotted line) are shown as bars. Mean ± SD from 3 independent experiments is plotted and p-values from Student’s t-test for relevant comparisons are shown. **C**. Experimental design for estimating the establishment probability of resistant mutants over wild type. Mixed strain cultures were set up with wild type in a large excess. Replicates (144 for each comparison) were set up in 96-well plates in drug-free or trimethoprim-supplemented (100 ng/mL) media and allowed to grow for 24 hours. An aliquot (1%) of the culture was passaged into media containing high trimethoprim (1 µg/mL i.e. MIC of wild type). Growth in high trimethoprim would be observed only in those cases where the resistant mutant had been enriched by >10-fold **D**. Scatter plot of OD values (at 600 nm) for all 144 competitions carried out to determine establishment propensity. Monocultures of wild type and competitions in drug-free media (gray) were used as controls to negate the effect of spontaneous resistant mutants emerging during the assay. Competitions in trimethoprim-supplemented media are plotted in ochre. Median OD values are shown as dashed lines. The fraction of replicates in which the mutant was enriched compared to wild type based on visible growth is indicated above the appropriate OD scatter.

In parallel, we also analysed the expression level of *folA* in these resistant strains. Loss of functional MgrB alone led to ∼2 fold higher levels of *folA* transcript than wild type (Figure 3B). Likewise, Isolate Tmp^R^-B also showed ∼2 fold higher *folA* transcript, while Tmp^R^-*folA*-Trp_30_Arg had comparable *folA* expression as wild type (Figure 3B). On the other hand, the Tmp^R^-*folA*-C_-35_T showed massive overproduction of *folA*, which was not detectably higher in strain Tmp^R^-A (Figure 3B). Thus, *mgrB*-mutations enhanced the expression of wild type or missense alleles of *folA*. However, the effect of *mgrB*-mutation was marginal if cis-regulatory changes in the *folA* promoter were present as the latter masked the effects of hyperactive PhoP. The strongly correlated trends between IC_50_ values and *folA* expression demonstrated that the molecular mechanism behind the observed epistasis was MgrB’s influence on *folA* expression levels.

Could this genetic interaction explain why mutations in *mgrB* facilitated fixation of resistant bacteria? To answer this question, we calculated the frequency at which resistant strains harboring different mutation combinations could establish over a large excess of a drug-sensitive competitor in the presence and absence of sub-MIC trimethoprim (100 ng/mL) (Figure 3C). Competitions (144 replicates for each strain combination) were set up with wild type *E. coli* at a starting ratio of 1:10^7^ in favour of wild type to mimic initial stages of evolution when a resistant mutant first emerges in an ancestrally sensitive population (Figure 3C). After 24 hours of competition, 0.5 % of the mixed culture was passaged into LB media containing trimethoprim at the MIC of wild type (1 µg/mL). Growth at this concentration of trimethoprim would indicate that the resistant mutant had been enriched >10-fold during the competition (Figure 3C). Wild type bacteria alone and competitions in the absence of trimethoprim were used as controls. Indeed, strains Tmp^R^-A and B had higher frequencies of establishment than Tmp^R^-*folA*-C_-35_T and Tmp^R^-*folA*-Trp_30_Arg demonstrating the impact of *mgrB* mutations as facilitators of resistance evolution (Figure 3D). Further, *mgrB*-mutation enhanced the frequency of establishment of the *folA*-Trp_30_Arg allele to a greater extent than *folA*-C_-35_T, in line with the differential epistatic effects between these mutations (Figure 3D). Thus, we concluded that *mgrB*-mutations facilitated the fixation of bacteria with mutations in *folA* under drug pressure, which could be explained mechanistically by epistasis between *mgrB* and *folA* loci.

### MgrB is retained by *E. coli* to prevent costs of PhoQP hyperactivation

Since loss of functional *mgrB* was frequent and beneficial under antibiotic pressure, we wondered why negative feedback may be needed in the PhoQP system in the first place. To address this question, we first assessed the evolutionary conservation of *mgrB* across PhoQP-expressing bacteria. The PhoQP system is restricted to the Order Enterobacterales [43]. Within this order, the *phoQ* gene was found in all bacterial families except Budviciaceae, indicating that PhoQP may have emerged after the divergence of Budviciaceae from the other 6 families (Figure 4A). The *mgrB* gene was present in representative members of 4 of the 6 families that harboured *phoQ*, namely Morganellaceae, Yersiniaceae, Pectobacteriaceae and Enterobacteriaceae, but not in Erwiniaceae and Hafniaceae (Figure 4A). Phylogenetic relationships between these families [44] suggested that Erwiniaceae and Hafniacieae may have independently lost *mgrB* during evolution. We next examined the distribution of *phoQ* and *mgrB* among members of family Enterobacteriaceae, to which *E. coli* belongs (Figure 4B). Among 61 query species examined by us, we found that most harboured both, *phoQ* and *mgrB*. A few exceptions were also identified, such as Izhakiella, Rosenbergiella, Limnobaculum and some species of Candidatus, which code for PhoQ but not MgrB (Figure 4B). Once again, their phylogenetic relationships suggested that these genera represented independent gene-loss events (Figure 4B). Taken together, these analyses showed that multiple independent events of loss of the *mgrB* gene have occurred during bacterial evolution. However, most extant bacterial species preserved a functional *mgrB* gene, indicating that negative feedback in PhoQP is dispensable but desirable.

**Figure 4.**
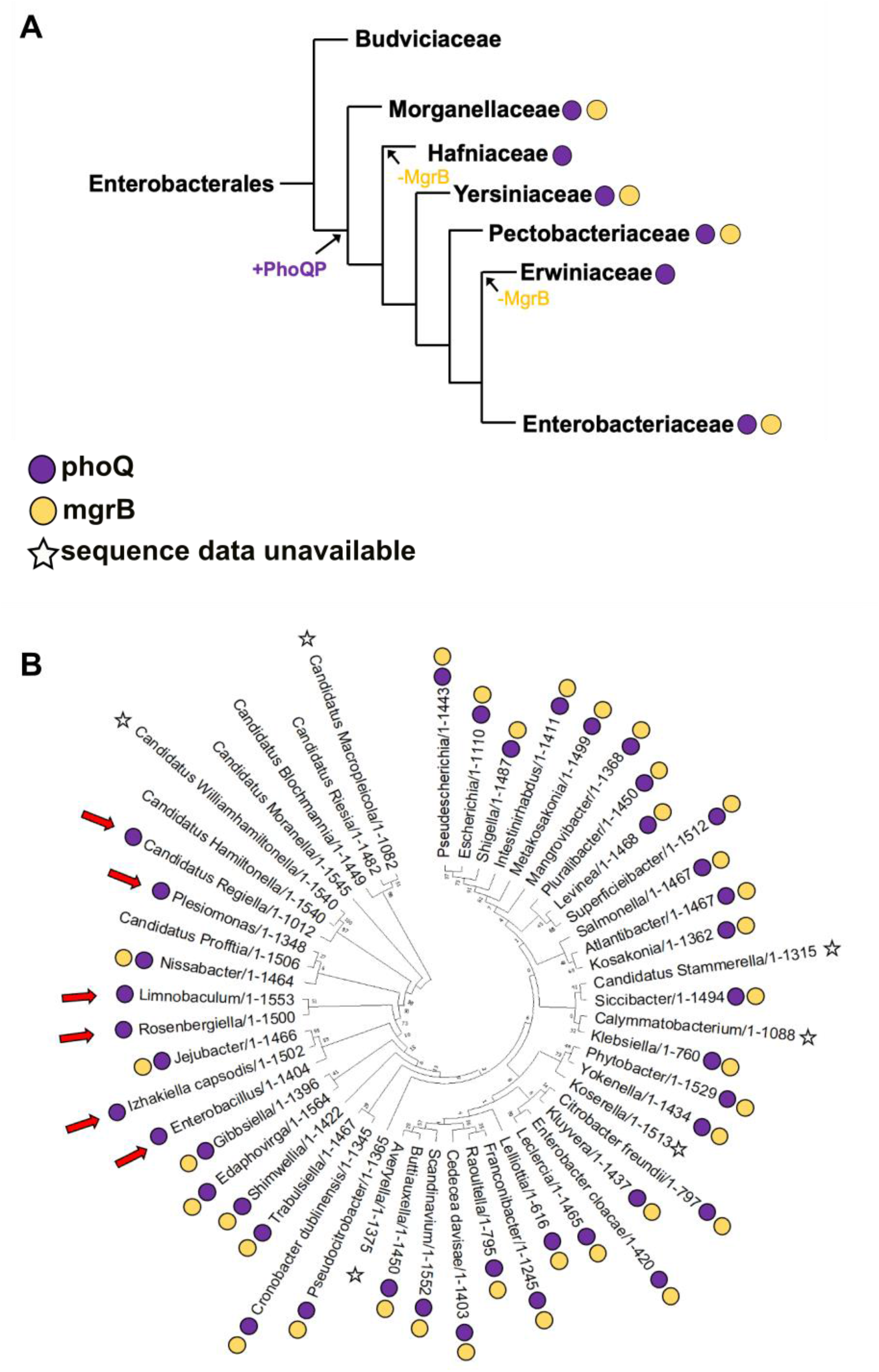
Phylogenetic distribution of *mgrB* and *phoQ* genes across bacteria. **A**. Representation of the phylogenetic relationships between 7 bacterial families that make up the Order Enterobacterales, taken from Adelou et al., 2016 [44]. The presence of *mgrB* and *phoQ* genes in representative members of these families are shown as yellow and purple circles respectively. Most probable events of gain of the PhoQP system and loss of MgrB are indicated. **B**. Maximum Likelihood phylogeny of 61 representative species belonging to Family Enterobacteriaceae constructed using 16S rRNA sequences is shown. The presence of *phoQ* and *mgrB* are indicated by purple and yellow circles next to each species. Lack of annotated genome data is indicated by a star. Possible events of *mgrB*-loss are shown as red arrows.

In line with this idea, there was no detectable difference in the growth characteristics of wild type *E. coli* and its isogenic *mgrB* knock-out strain under standard laboratory conditions (Figure 5A). However, when *E. coli ΔmgrB* was directly competed against *ΔlacZ-*marked wild type, a significant fitness cost (w_ΔmgrB_ = 0.83±0.03) was observed (Figure 5B, C). The fitness cost of *mgrB*-deficiency was due to hyperactivation of PhoQP since addition of high Mg^2+^ to growth media or deletion of *phoP/Q* genes restored relative fitness to wild type levels (Figure 5B, C). Our earlier work has shown that loss-of-function mutations in *rpoS* that occurred spontaneously during long term evolution in trimethoprim improved the fitness of *mgrB*-deficient *E. coli* in the drug-supplemented media without enhancing IC_50_ [33]. Since RpoS is itself an indirect target of PhoQP signaling and known to be overproduced in *mgrB*-knock out bacteria [45], we asked whether loss of *rpoS* could compensate for the fitness costs of *mgrB*-deficiency in drug-free media as well. This was indeed the case and deletion of *rpoS* restored the relative fitness of an *mgrB*-knockout strain (Figure 5B, D). Like *rpoS*, deletion of *iraM*, which links *rpoS* to PhoQP signaling [45], also rescued the fitness cost of *mgrB*-deficiency (Figure 5B, D). Thus, we concluded that the costs of *mgrB*-deficiency could be mechanistically traced to hyperactivation of PhoQP and over-production of RpoS, explaining why negative feedback may be needed in the PhoQP system.

**Figure 5.**
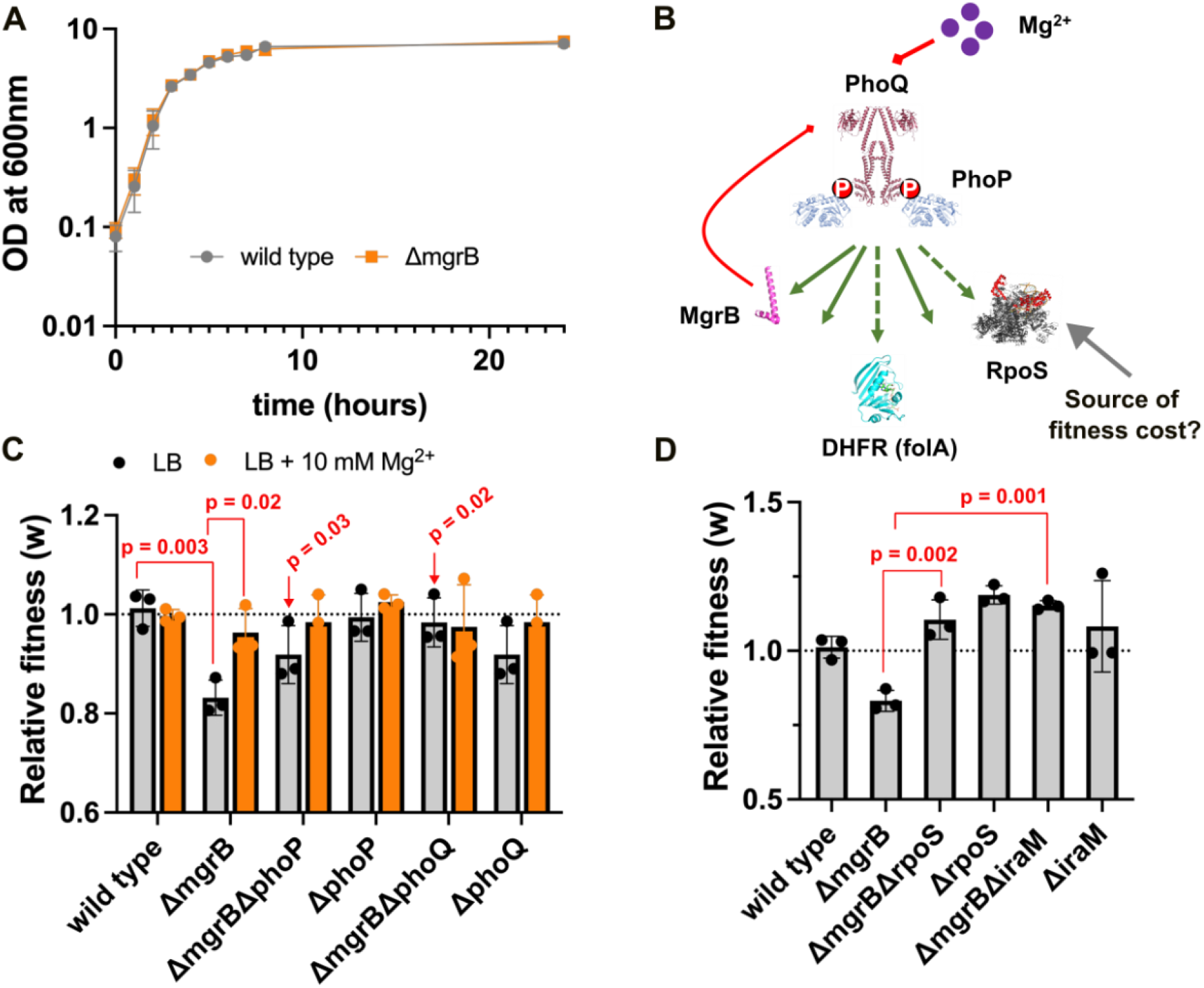
Loss of negative feedback is costly for *E. coli* due to PhoQP hyperactivity. **A**. Growth curves of wild type *E. coli* and an isogenic *mgrB*-knockout strain *(ΔmgrB*) under standard growth conditions in LB. Growth was measured using OD at 600 nm. Mean ± SD from 3 independent experiments is plotted. **B**. Diagrammatic representation of the PhoQP signaling pathway (not to scale). Direct and indirect positive regulation by PhoP is shown as solid and dashed green arrows respectively. Inhibitory interactions are shown in red. AlphaFold models/crystal structures are used to represent proteins in the pathway that are relevant to this study. Source of the fitness cost of *mgrB*-deficiency based on our study is shown. **C and D**. Relative fitness (w) of *E. coli* wild type and indicated mutant strains compared to a *lacZ*-knock out reference strain calculated in LB (gray bars; low Mg^2+^) or LB + 10 mM MgSO_4_ (orange bars). Mean value from three independent experiments are plotted as bars, and individual values are shown. Error bars represent standard deviation. A value of 1 represents no change in fitness compared to the reference strain and is shown as a dotted line. Statistical significance was tested using a Student’s t-test (p-values for relevant comparisons are shown). Double mutants were compared with *ΔmgrB* for statistically significant differences.

### Pervasive gene dysregulation in *mgrB*-deficient *E. coli* is rectified by mutations in *rpoS*

We next sought to understand the molecular mechanistic basis for the cost of *mgrB*-deficiency and the compensatory role of *rpoS*-mutations. To do this, we compared the whole transcriptomes of trimethoprim-resistant isolates from early and late time points of our long-term evolution lines using RNA-seq. The isolates Tmp^R^-A and Tmp^R^-B were chosen from early time points since they both harboured loss-of-function mutations at the *mgrB* locus and had wild type *rpoS* (Figure 6A). Since both isolates also harboured other mutations, comparing the transcriptomes of Tmp^R^-A and Tmp^R^-B would help to identify *mgrB*-specific gene regulatory effects. The isolate Tmp^R^-C was chosen from a later time point as it harboured the same mutation in *mgrB* as Tmp^R^-B and an inactivating mutation in *rpoS* (Figure 6A). All three isolates also harboured mutations in the *folA* gene (Tmp^R^-B, C) or its promoter (Tmp^R^-A) (Figure 6A).

**Figure 6.**
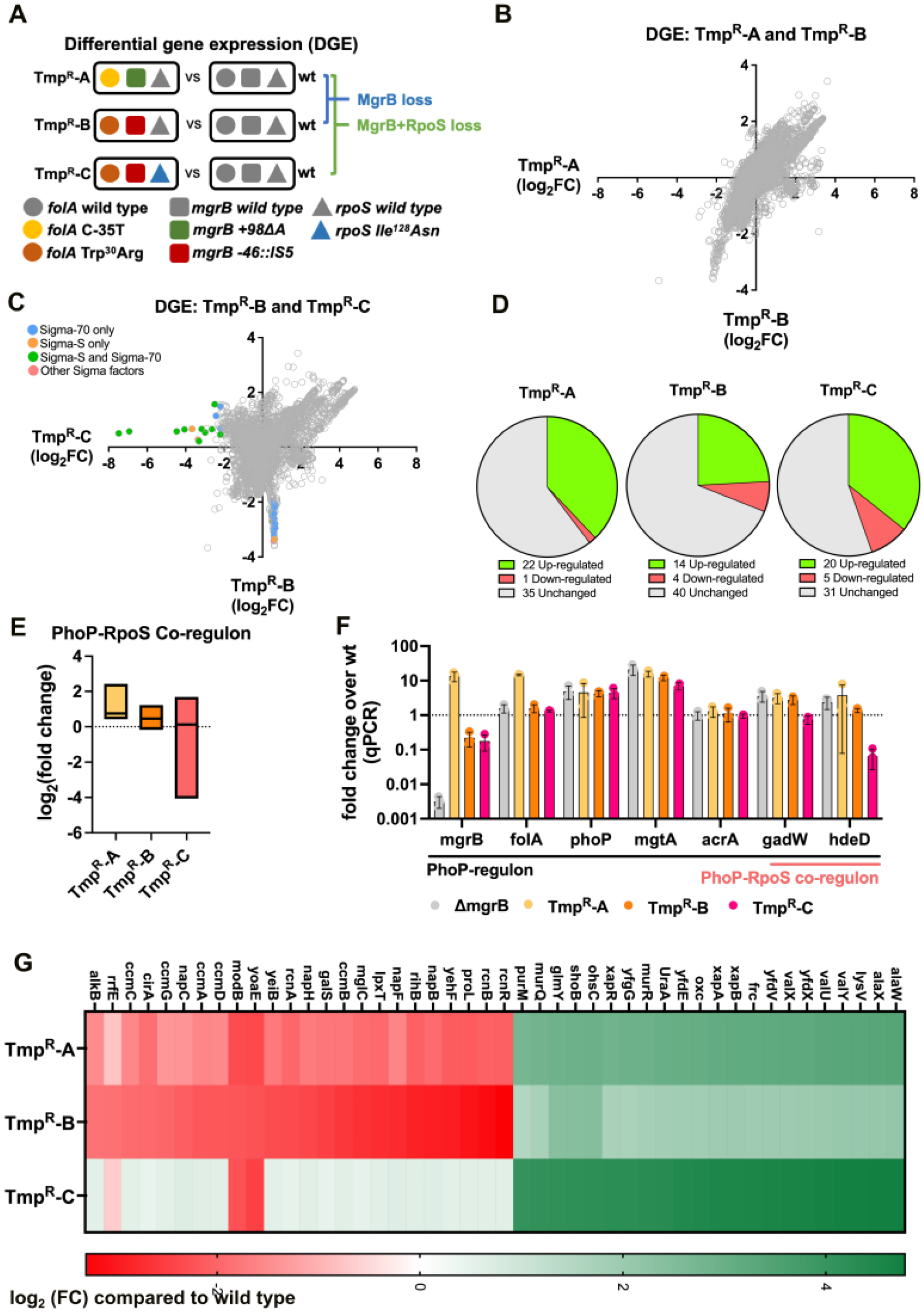
Transcriptional changes in trimethoprim resistant isolates explain the compensatory role of *rpoS* mutations. **A**. Diagrammatic representation of the genotypes of strains used for RNA-sequencing based transcriptomics. Circles represent *folA*, squares represent *mgrB* and triangles represent *rpoS*. Gray symbols represent wild type alleles, while coloured symbols represent mutant alleles as shown in the key. Comparisons between strains reveal the effects of *mgrB* and *mgrB*+*rpoS* mutations on the global transcriptome of *E. coli* as shown. **B**. Comparison of differential gene expression (DGE) between trimethoprim resistant strains Tmp^R^-A and Tmp^R^-B. For each strain DGE was estimated as log_2_ of fold change compared to wild type (log_2_FC). Each point on the scatter plot represents expression level of a single gene. **C**. Comparison of differential gene expression (DGE) between trimethoprim resistant strains Tmp^R^-B and Tmp^R^-C. For each strain DGE was estimated as log_2_ of fold change compared to wild type (log_2_FC). Each point on the scatter plot represents expression level of a single gene. For genes showing greater than 4-fold down-regulation in Tmp^R^-B or Tmp^R^-C compared to wild type, data points are colored by the sigma factor responsible for transcription as indicated based on data available on Ecocyc database [46]. **D**. Expression level of 58 genes of the PhoP regulon (gene list obtained from RegulonDB [47, 48]) in Tmp^R^-A, B and C strains represented as pie-charts. Genes with at least 2-fold higher or lower expression level compared to wild type were classified as “up-regulated” or “down-regulated” respectively. **E**. Expression level of 14 genes co-regulated by PhoP and RpoS in Tmp^R^-A, B and C strains shown as a box plot (min to max). The median expression level is shown by the line. **F**. Validation of RNA-sequencing based transcriptomics using quantitative RT-PCR (qPCR) for selected genes. Mean ± SD of fold change compared to the wild type from three independent biological replicates are plotted. No change in expression level compared to wild type corresponds to a value of ‘1’ and is shown by a dotted line. Gene names are indicated on the X-axis. Different colours represent different strains from which measurements were made as shown in the key below the graph. **G**. Heat map showing expression level of RpoD-target genes in Tmp^R^-A, B and C strains based on RNA-seq. For each strain, expression levels were compared to wild type and log_2_(FC) is represented on a continuous red-green colour scale as shown.

At the global level, the differential gene expression profiles of Tmp^R^-A and Tmp^R^-B were very similar and strongly correlated (linear correlation coefficient R^2^ = 0.59) (Figure 6B). On the other hand, Tmp^R^-B and Tmp^R^-C showed significant differences in their transcriptomes (linear correlation coefficient R^2^ = 0.19) (Figure 6C). Next, we turned our attention specifically to genes of the PhoP-regulon. A total of 58 genes are known direct targets of the PhoQP system in *E. coli* [46-48]. We noted that despite the presence of *mgrB*-mutations, majority of the PhoP-regulon remained unaltered in all 3 isolates (Figure 6D, Supplementary File 3). For instance, the levels of PhoP-targets such as the *acrAB* efflux pump and the *glg* glycogen metabolism operon were unaffected by the loss of *mgrB*. These genes are regulated by other transcription factors in addition to PhoP, including master regulators such as CRP [46-48], and hence may be less sensitive to the activation status of PhoP. Hyperactivation of PhoQP was reflected most dramatically in the overexpression of genes that are solely regulated by PhoP, such as the *phoQP* operon itself or the magnesium transporter *mgtA* (Figure 6D, F, Supplementary File 3).

A subset of PhoP-targets is also regulated by the RpoS sigma factor [46-48]. Targets of PhoP and RpoS include genes that form the ‘acid-resistance regulon’ such as the ‘*gad*’ and ‘*hde*’ operons [45]. In Tmp^R^-A and B most of the PhoP-RpoS co-regulon showed mild to high over-expression compared to wild type (Figure 6E, Supplementary File 3). However, in Tmp^R^-C, several of these genes were significantly down-regulated relative to wild type, indicating that mutation in *rpoS* overrides the effects of PhoQP hyperactivation and compensates for the hyperactivity of PhoP (Figure 6E, Supplementary File 3). We confirmed these findings using qPCR for representative genes and found the results to be in line with the RNA-seq transcriptomics data (Figure 6F).

Curiously, we noticed that there were several differentially expressed genes between isolates that are not direct targets of PhoP or RpoS (Supplementary File 3). Particularly striking among them were genes that were significantly down-regulated in Tmp^R^-A and Tmp^R^-B but restored to wild type levels in Tmp^R^-C (Figure 6C, Supplementary File 3). A closer look at this sub-set of genes revealed that a majority of them were transcribed in an RpoD-dependent (i.e. Sigma 70-dependent) manner and were involved in growth and metabolism such as *lpxT* (lipid A metabolism), *ccmA-D* (cytochrome maturation) and *rcnA-B* (metal ion homeostasis) (Figure 6C, G, Supplementary File 3). Similarly, several RpoD-regulated tRNA genes were also expressed to a higher level in Tmp^R^-C, compared to Tmp^R^-A and Tmp^R^-B (Figure 6G, Supplementary File 3). It is well-established that RpoS and RpoD compete for the same binding site on RNA polymerase and competition between these two sigma factors dictates whether *E. coli* expresses genes required for growth and division (RpoD-regulated) or stress-response and stationary phase (RpoS-regulated) [49]. The dysregulation of RpoD-target genes in Tmp^R^-A and B thus suggested that precocious RpoS activation due to *mgrB*-deficiency tilted the balance in favour of the RpoS-transcriptional program, which was compensated at later stages in evolution by mutations in *rpoS*. Based on these results, we concluded that loss of *mgrB* led to pervasive perturbation of gene expression beyond the PhoP-regulon, its primary target, to secondary and tertiary effects on the expression of RpoS- and RpoD-regulated genes respectively. Mutations in *rpoS* restored several of these secondary and tertiary gene regulatory effects.

### RpoS-RpoD imbalance explains the fitness costs of MgrB-deficiency in *E. coli* and justifies the need for negative feedback in the PhoQP system

Based on the above result, we hypothesized that the cost of *mgrB*-loss may arise either due to overproduction of RpoS-regulated genes or the repression of RpoD-regulated genes. To test which of these two possibilities was true, we first generated knockouts of the *hdeD, gadW* and *gadE* genes (co-targets of RpoS and PhoP) in an *mgrB*-deficient background and asked whether these gene deletions could rescue the fitness cost of the *mgrB* knockout. These specific genes were selected since they are master regulators of the acid-resistance regulon and were up-regulated in Tmp^R^-A and B, but significantly repressed in Tmp^R^-C. However, the fitness of the *mgrB* knockout was unaffected by deletion of these genes (Figure 7A) ruling them out as the source of the observed fitness cost. Next, to test whether the imbalance between RpoS and RpoD could explain the costs of *mgrB*-deficiency, we deleted the *rsd* or *crl* genes from the *mgrB*-knockout and measured relative fitness. Rsd is an inhibitor of RpoD [50, 51], while Crl is potentiator of RpoS [52], and loss of either protein would tilt the balance in favour of RpoD-regulated transcription. Indeed, deletion of *rsd* or *crl* rescued the costs of the *mgrB*-knockout strain (Figure 7A). Finally, we traced how the competitive ability of *E. coli ΔmgrB* changed with growth phase. We found that in the logarithmic phase, when RpoD is known to be higher, there was no detectable difference between *mgrB*-deficient and wild type bacteria in a mixed culture. However, the competitive disadvantage of *E. coli ΔmgrB* became evident at the onset of stationary phase when the shift from RpoD to RpoS-mediated transcription is known to occur (Figure 7B). This effect too was compensated by deletion of *rpoS* (Figure 7B). These results confirmed that pervasive transcriptional dysregulation of gene expression due to hyperactive PhoQP explained the costs of *mgrB* deficiency and provided a mechanistic explanation for why negative feedback is retained by this two-component system during evolution.

**Figure 7.**
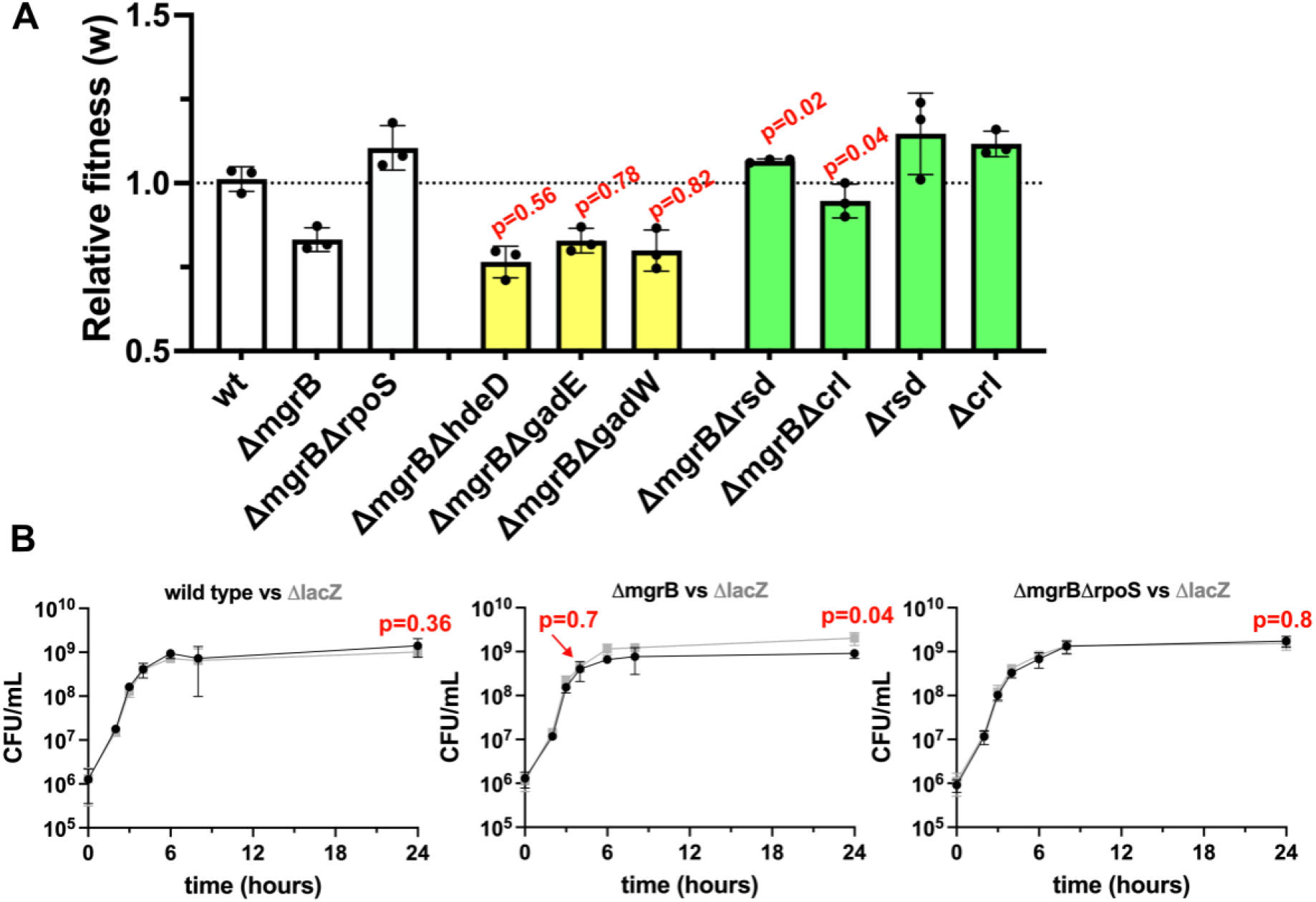
Loss of RpoS-RpoD balance explains the fitness costs of *mgrB*-deficiency. **A**. Relative fitness (w) of *E. coli* wild type and indicated gene knockout strains compared to an isogenic *lacZ*-knock out reference strain in antibiotic-free media. Mean value from three independent experiments are plotted as bars. Error bars represent standard deviation. A value of ‘1’ indicates no change in fitness compared to the reference strain and is shown as a dotted line. Yellow bars represent strains that were used to test the contribution of RpoS-targets to the costs of *mgrB*-deficiency, while green bars represent strains that were used to test the contribution of RpoS-RpoD imbalance to the costs of *mgrB*-deficiency. Statistical significance between the relative fitness of double mutants and *ΔmgrB* was tested using a Student’s t-test (p-values shown above relevant bars). **B**. Growth of wild type, Δ*mgrB* or Δ*mgrB*Δ*rpoS* (black) in competition with an isogenic Δ*lacZ* strain (gray) over time expressed as CFUs/mL. Mean ± SD from three independent biological replicates is plotted. Student’s t-test was used to compare statistical significance of the differences in bacterial densities of the competing strains (p-values are shown next to relevant data points).

## Discussion

Loss of functional MgrB occurs spontaneously under antibiotic pressure and is associated with resistance to trimethoprim in *E. coli* [33, 38] and colistin in *K. pneumoniae* [39]. Being a negative feedback regulator of the PhoQP two-component system, absence of MgrB hyperactivates PhoP and leads to overexpression of several of its targets [37, 45]. In this study, we have used the PhoQP/MgrB two-component system from *E. coli* to ask what the contribution of gene regulatory evolution is to antimicrobial resistance in bacteria. Our results demonstrate that loss of checks and balances such as negative feedback in gene regulatory pathways can be adaptive for bacteria under antibiotic pressure. Further, deregulating gene expression has the potential to amplify the phenotypes of other resistance-conferring mutations and hence facilitate the evolution of high level antibiotic resistance. Importantly, we have also shown that deregulation of signalling pathways has pervasive effects on the wider gene regulatory network of cells, which are maladaptive. As a result, it is accompanied by a fitness cost that must be compensated through additional evolutionary adaptation (Figure 8).

**Figure 8.**
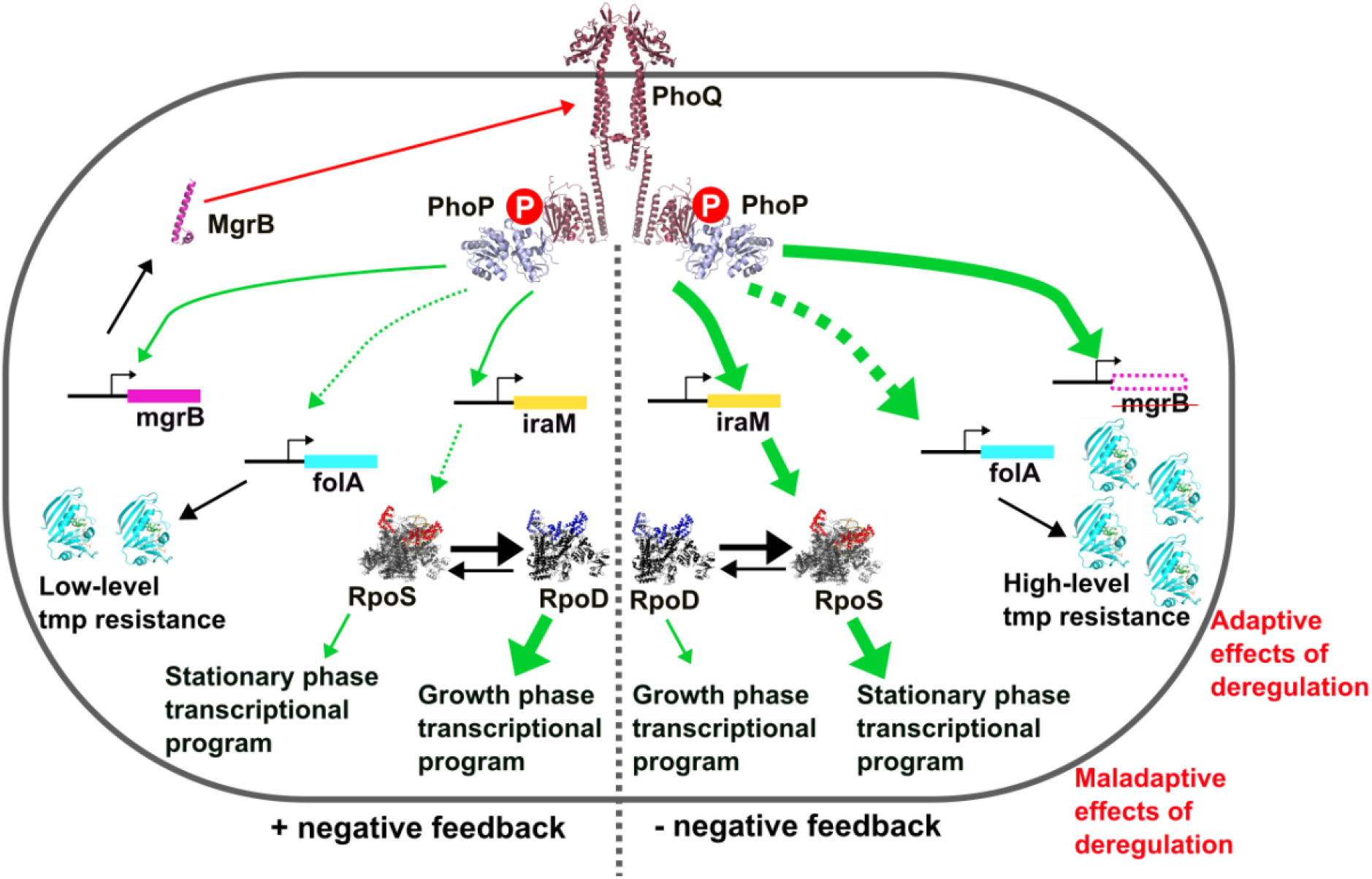
A gene regulatory model for the adaptive and maladaptive effects of deregulation in the PhoQP pathway. Diagrammatic representation of the proposed model for why deregulation of PhoQP due to loss of negative feedback regulator MgrB produces both adaptive and maladaptive gene regulatory changes. Left and right sides of the figure represent scenarios where negative feedback is retained or lost by the PhoQP system respectively. Direct and indirect effects are represented by solid and discontinuous arrows respectively. Positive regulation is represented by green arrows while negative regulation is represented by red arrows. Arrow thickness indicates strength of effect. Genes and their promoters are represented by boxes, while proteins are shown as structural models (Alphafold; https://alphafold.ebi.ac.uk/). The figure is not drawn to scale.

Increased expression of drug-targets, most commonly due to mutations in gene promoters or transcription factors, is reported in many antibiotic-resistant bacterial pathogens. For instance, mutations in the promoter of the *pbp4* gene in beta-lactam resistant *Staphylococcus aureus* [53], missense mutations in *mgrB* or *phoQ* in colistin-resistant *Klebsiella pneumoniae* [54] and *inhA* promoter mutations in isoniazid-resistant *Mycobacterium tuberculosis* [55] are all frequently-encountered, clinically-relevant mechanisms of antibiotic resistance. The main role of regulatory mutations is assumed to be to enhance bacterial fitness by increasing expression of the drug target. In our study, loss of functional MgrB, and associated overexpression of *folA* did indeed confer a fitness advantage to *E. coli* challenged with trimethoprim. By itself, this benefit was incremental and would normally be considered clinically irrelevant. However, *mgrB*-loss contributed significantly to the evolution of high-level trimethoprim resistance by synergistic epistasis with missense mutations in *folA*. Consequently, *mgrB*-mutations were the most frequent genetic change in laboratory-evolved trimethoprim resistant bacteria despite the small stand-alone impact on drug IC_50_/MIC. Thus, our study throws light on another, perhaps more significant role of regulatory mutations as facilitators of antimicrobial resistance evolution through epistasis. Phenomenologically similar ideas have been proposed by a few other studies, though in these cases the mechanistic bases were unclear. For instance, antibiotic-induced alterations in global gene expression promoted the fixation of resistance-conferring mutations in *E. coli* challenged with amoxicillin and tetracycline [56]. Similarly, expression-modifying mechanisms have been shown to serve as “latent defences” that bacteria can exploit in order to adapt to antibiotics [57]. Taken together, the role played by regulatory mutations in facilitating the evolution of antimicrobial resistance is likely to be more wide-spread than understood so far and requires greater attention.

Based on our previous work, as well as work from other groups, we explain the mechanism of epistasis between *mgrB* and *folA* as follows. Missense mutations in DHFR can structurally destabilise the protein and reduce its steady state levels [58, 59]. Hyperactive PhoP is likely to compensate for this effect by increasing the transcription of mutant DHFR in the bacterial cell, resulting in high levels of drug resistance. Interestingly, we have shown earlier that loss of Lon protease activity also shows synergy with unstable drug-resistant DHFR mutants by increasing their *in vivo* half-life [60]. Thus, increasing expression level of mutant DHFRs, regardless of mechanism, can facilitate high-level resistance evolution. This mechanism is consistent with our observation that coding mutations in DHFR synergize with *mgrB*-mutations, while mutations in the *folA* promoter don’t do so. It is important to note here that the number of possible resistance-conferring coding-region mutations in *folA* is much higher than promoter mutations [33, 61-63]. Thus, by-and-large *mgrB* is more likely to synergistically enhance the resistance level of *folA*-mutant bacteria than not.

We propose that characterising gene regulatory evolution in drug-resistant bacteria may help to identify new points of intervention for AMR pathogens. Our study shows that “switching-off” PhoQP signalling sensitized resistant *E. coli* to trimethoprim as well as retarded resistance evolution. Sensitization strategies such as this one are potential solutions to the current AMR crisis. A few approaches to achieve sensitization to antibiotics that have been explored so far have used adjuvant molecules that enhance the concentration of antibiotics in bacterial cells [64]. For instance, efflux pump inhibitors like verapamil have been explored as a possible mode of re-sensitising multi-drug resistant bacteria [65]. Similarly, cell wall perturbing polymers can also potentiate the activity of some antibiotics by enhancing their penetration into bacterial cells [65, 66]. Our work shows that modifiers of two-component system-regulated gene expression may serve as an additional or alternative sensitization strategy. Though our results have been limited to trimethoprim resistant *E. coli*, several two-component systems are associated with resistance to antibiotics in different bacteria, often due to direct activation by the antibiotic [27]. Thus, this strategy for sensitization is likely to have wide application. Indeed, there are also potential advantages to targeting two-component sensor kinases for therapeutic purposes. Since they are not found in human cells, pharmacologically inhibiting two-component may produce fewer off-target effects in the host. Further, being cell envelope proteins with known ligands, inhibitor design may be facilitated. Finally, modulation of bacterial gene expression has not yet been targeted therapeutically and hence is unlikely to be confounded by pre-existing resistant mutants in clinical strains. These interventions may also serve the purpose of “evolution-proofing” of new antibiotics since they are likely to slow down the evolution of resistance as shown by us in this study and could be thought of as prospectively-used adjuvants.

Beyond the evolution of drug-resistance, our study contributes to the understanding of how gene regulatory networks evolve, particularly in the context of the emergence and maintenance of negative feedback. Mathematical modelling coupled with experiments using natural or synthetic genetic circuits have shown that negative feedback can reduce noise in gene regulatory networks [3, 5, 10, 11, 67]. Despite this knowledge, the importance of negative feedback from an evolutionary perspective remains poorly investigated. The only empirically validated basis for the evolution of negative feedback is to enhance mutational tolerance. Marciano et al. used the LexA transcription factor of *E. coli* to demonstrate that negative feedback canalises phenotypes and provides greater tolerance to mutational perturbation [68]. Similar observations were made for the Rox1 protein from *Saccharomyces cerevisiae* where negative feedback stabilised gene expression levels and enhanced mutational robustness [69]. For the PhoQP system, it has been speculated that MgrB may have evolved to optimise the signalling output of PhoQP in environments with varying Mg^2+^ concentrations [34, 36, 37]. Our study provides experimental evidence for an alternative explanation for the evolution of negative feedback in PhoQP, i.e., to prevent pervasive dysregulation of the larger bacterial genetic network. In other words, we propose that negative feedback serves to insulate pathways that can potentially cross-activate, like PhoQP and RpoS transcriptional networks.

We note that wider application of the above idea to other bacteria is contingent on the pervasive gene regulatory effects of PhoQP in different bacterial species. Indeed, the PhoQP regulon has been worked out in a few species other than *E. coli*. We find that in all characterised systems, like in *E. coli*, PhoQP activation leads to gene expression changes that extend beyond the PhoP regulon. For example, in *Pectobacterium versatile* and *Yersinia pestis*, genome-wide transcriptomics and ChIP-seq have revealed that PhoP influences the expression of several genes beyond its direct targets, though the mechanisms aren’t yet clear [70-72]. For *E. coli*, our study shows that RpoS overproduction and exclusion of RpoD from RNA polymerase are responsible for activating secondary and tertiary transcriptional programs in response to PhoQP hyperactivity. Interestingly, while Pectobacterium and Yersinia code for RpoS, they lack IraM-like proteins and hence in these species PhoQP may produce pervasive dysregulation of gene expression by other mechanisms [43]. Regardless, a broad gene-regulatory influence of the PhoQP pathway seems to be consistent across bacteria, supporting the idea that a key selection pressure for the maintenance of MgrB may be to limit cross-activation of other transcriptional programs by PhoP. It is important to note here that the requirement for negative feedback is highly contextual, at least in pathways like PhoQP that directly respond to the extracellular medium. Indeed, deregulation of PhoQP by loss of MgrB is itself beneficial in many environments such as acid stress and antibiotics [33, 39, 45, 54]. The multiple instances of loss of *mgrB* across bacterial phylogenies observed by us in the present study may reflect different selection pressures driving evolution of the PhoQP system in different bacteria. Similarly, cross-activation of other regulatory pathways, though costly under the conditions tested by us, may have advantages in more complex environments. For the PhoQP/MgrB system from *K. pneumoniae* this idea has been recently proposed. Bray et al. (2021) [73] demonstrated that loss of MgrB in colistin resistant-*K. pneumoniae* is accompanied by compromised gastrointestinal colonisation efficacy. Like our results, this study too reported the overproduction of RpoS in *mgrB*-deficient *K. pneumoniae*. However, the authors reported that RpoS overproduction was beneficial in a model for pathogen transmission [73]. Thus, life-history and growth context are both likely to have a strong influence on how deregulation of gene expression translates to organismal fitness.

Curiously, the evolution of negative feedback proteins in two-component signalling pathways appears to be the exception rather than the rule. In *E. coli* only 2 systems, i.e. PhoQP and CpxAR have known negative feedback regulator proteins [74]. Our results may throw light on why this is the case. Based on the idea that pervasive gene dysregulation drives the evolution of negative feedback, we propose that the following criteria necessitate the evolution of negative feedback in two-component pathways. Firstly, the system must have a large regulon which increases the chance of its activity translating to changes in organismal fitness across environments. Secondly, the system should have a positive feedback loop, i.e., it should activate its own expression. This is known to be true for the PhoQP system and results in rapid amplification of signalling after activation. Finally, the regulon of the two-component system should be connected to other global regulatory networks, such as RpoS in the case of PhoQP. For *E. coli*, only 3 two-component systems satisfy all these criteria, namely PhoQP, CpxAR and ArcAB, of which 2 have known negative feedback regulators (Figure 9). In the case of ArcAB, we are not aware of negative feedback systems, however we believe that it may be reasonable to look for them in the future. We cannot rule out the role of regulatory RNAs here, several of which are known to be activated by two-component signalling [75]. Further investigation would be needed to test whether their contribution is similar to that of MgrB.

**Figure 9.**
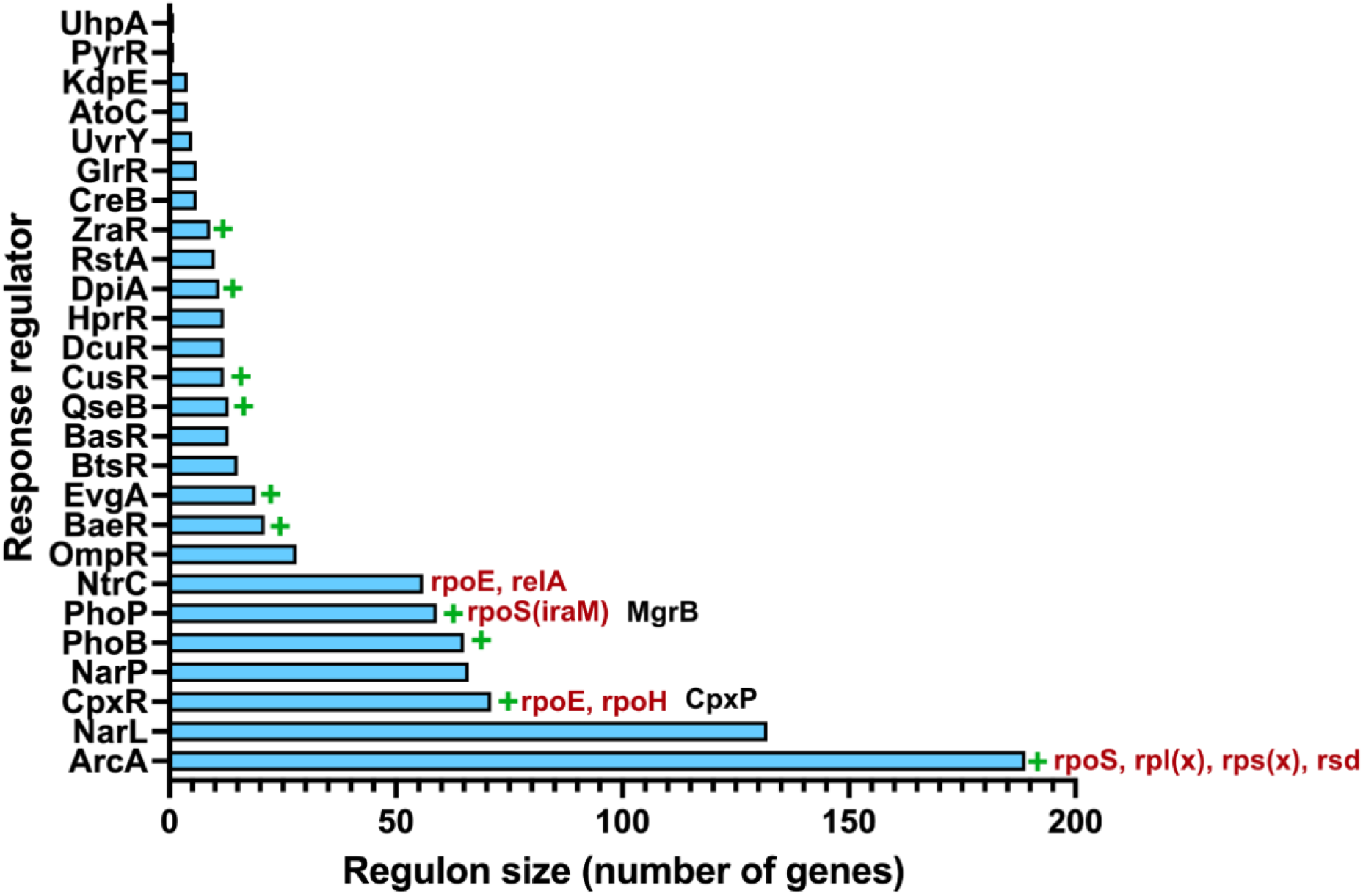
Regulon size and feedback in two-component systems of *E. coli*. Response regulators of 26 two-component systems from *E. coli* K-12 MG1655 ranked by their regulon sizes. Green plus sign (+) indicates that the two-component system is known to have positive feedback, i.e. stimulate its own transcription. Global regulators that these systems are known to cross-activate are in crimson next to respective response regulators, while known negative feedback regulatory proteins are in black. All information collated from Ecocyc [46].

In conclusion, despite extensive mechanistic insights into the functioning of gene regulatory networks, there is relatively less known about how they evolve and are rewired in response to environmental perturbation. Using the PhoQP-MgrB system, we have analysed the immediate and distal effects of deregulation by loss of negative feedback on the cellular transcriptional network. By linking these changes with organismal fitness across relevant environments we have shown how evolution in a gene regulatory network proceeds in response to selection and can drive and modulate evolutionary adaptation.

## Materials and Methods

### Bacterial strains and culture conditions

*E. coli* wild type and its mutants were cultured in Luria-Bertani Broth (LB) or on Luria-Bertani Agar (LA plates). Media were supplemented with trimethoprim and MgSO_4_ at required concentrations as needed. Kanamycin or chloramphenicol for selection of genetically manipulated strains were added at 30 µg/mL each as needed.

The strains used in this study are shown in Table 1.

**Table 1.**
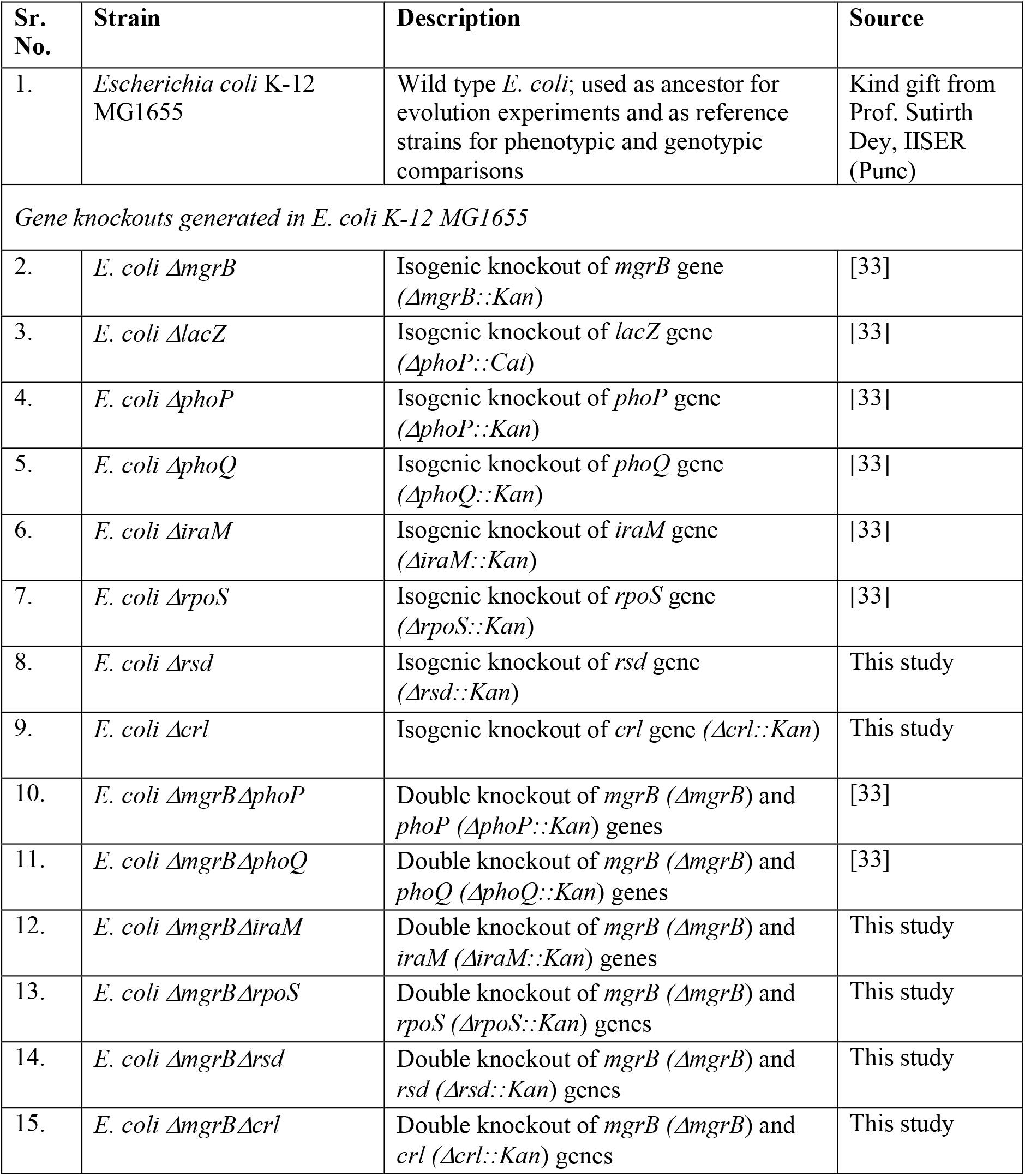
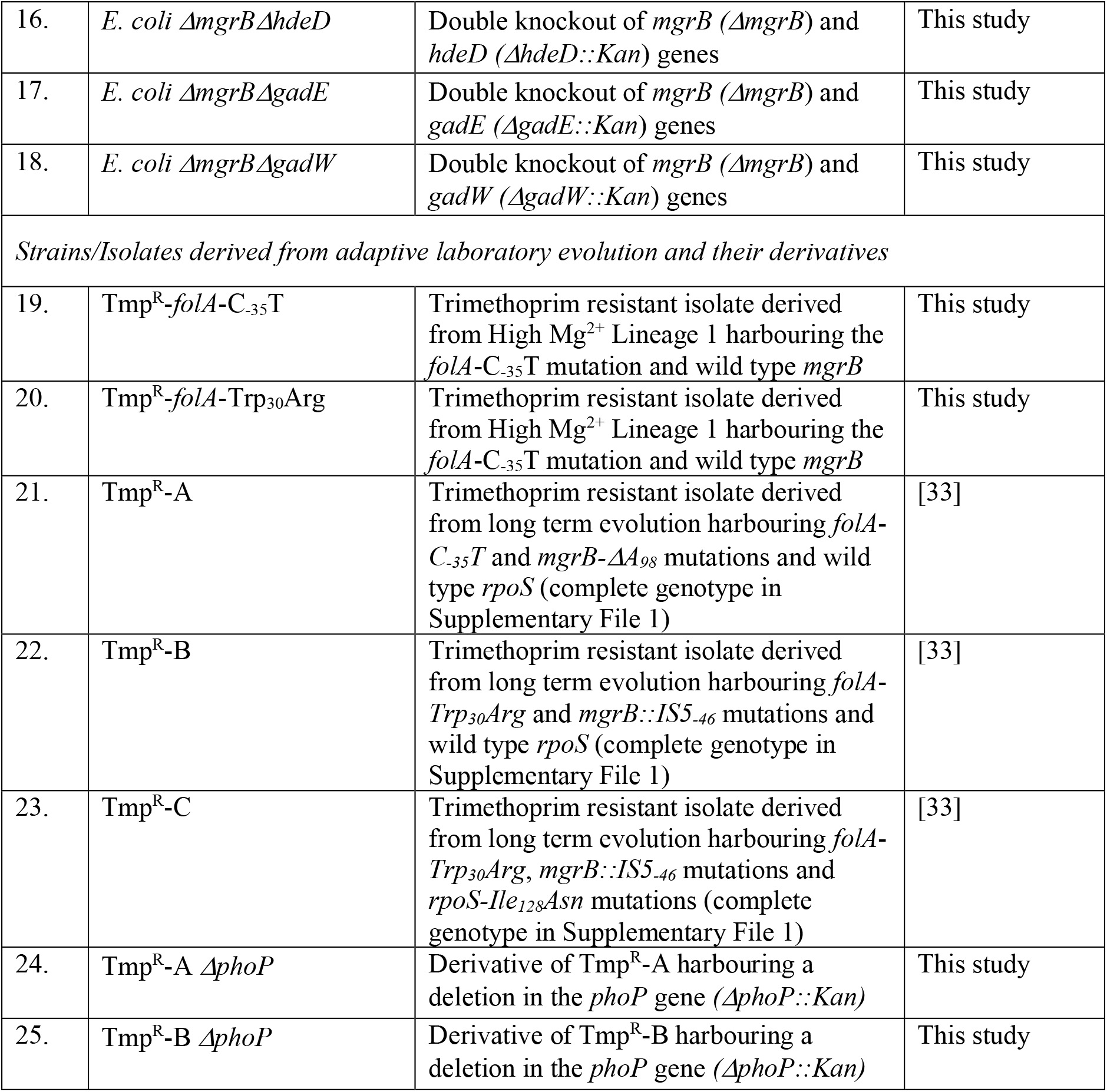
List of strains used in this study.

### Relative fitness measurements

Relative fitness (w) of various mutant strains of *E. coli* was measured by direct competition with an *E. coli ΔlacZ* strain under appropriate growth conditions. Neutrality of the *ΔlacZ* genetic marker was established by competitions with unmarked wild type *E. coli* under every growth condition tested. The detailed methodology followed is described in Patel and Matange (2021) [33].

### Monoculture and mixed culture growth curves

Bacterial strains to be characterised were initially grown overnight to saturation. From saturated cultures, bacteria were passaged (0.1%) into 5 mL LB broth and grown at 37°C, with shaking at 180 rpm. For competitive growth curves, 2.5 μL each of the competing strains were inoculated in 5 mL LB.

Aliquots were taken for measuring bacterial growth periodically until 20-24 hours of growth. Bacterial growth was monitored using optical density (OD) at 600 nm for monocultures and by viable counts (CFU/mL) for mixed culture. For viable counts, aliquots of bacterial cultures were serially diluted and plated on LA supplemented with IPTG (50 µg/mL) and X-Gal (50 µg/mL). Plates were incubated for 18-24 hours at 37 ºC and blue and white colonies were counted.

### Laboratory evolution of trimethoprim resistance

The detailed methodology for laboratory evolution of trimethoprim resistance is described in Patel and Matange (2021) [33] and Vinchhi et al. (2023) [76]. Briefly, bacterial populations were grown in trimethoprim supplemented LB (low Mg^2+^) or LB + 10 mM MgSO_4_ as required for 15-18 hours before passaging (1%) into fresh media. Bacteria were passaged for ∼210 generations (6-7 generations per growth cycle) and aliquots periodically were frozen at -80 ºC for further analyses. Trimethoprim was used at a concentration of 100 ng/mL which corresponds to ∼MIC/9.

### Genome sequencing of laboratory-evolved trimethoprim resistant isolates

Genome sequencing and variant calling for evolved bacteria was carried out as described in Patel and Matange (2021) [33] and Vinchhi et al. (2023) [76]. Sequencing services were provided by Eurofins, India.

### Genetic manipulations and gene knockout in *E. coli*

All gene knockouts were generated using P1 transduction by moving kanamycin-resistance marked gene deletions from donor strains taken from the Keio Collection [77, 78] into appropriate recipient strains. Knockouts were confirmed using gene specific PCR on genomic DNA extracted from transductants. Detailed methodology is described in Patel and Matange (2021) [33].

### Measuring trimethoprim resistance

Trimethoprim resistance of *E. coli* strains or trimethoprim-resistant isolates was measured using a broth-dilution assay. Briefly, appropriate strains of *E. coli* were grown to saturation and then inoculated into serially diluted trimethoprim-containing media. Growth was monitored after 18-20 hours of growth and IC_50_ values were determined by fitting experimental data to a variable-slope, 4 parameter model using Graphpad Prism (version 9.1.4). Detailed methodology is described in Patel and Matange (2021) [33] and Vinchhi et al. (2023) [76].

### Establishment propensity of trimethoprim-resistant isolates at sub-MIC drug pressures

To estimate the propensity of establishment of trimethoprim-resistant strains, their ability to out-compete a large excess of wild type *E. coli* under sub-MIC trimethoprim selection was evaluated. To set up the primary competitions, trimethoprim-evolved mutant isolate and wild-type *E. coli* were mixed (1:10^7^), and the mixture was diluted 100 times in a final volume of 15 mL LB broth supplemented with 100 ng/mL trimethoprim. This diluted culture was then dispensed into the wells of a sterile 96-well polystyrene plates (200 µL per well), and incubated at 37°C for 24 hours, with shaking at 180 rpm. Next, 1 μL from each well was passaged into 200 μL of LB broth containing 1 μg/mL trimethoprim (i.e. MIC of wild type [33]) in a fresh 96-well plate. This plate was incubated at 37°C for 24 hours with shaking at 180 rpm. Only those wells in which mutant bacteria were enriched by >10-fold would show turbidity. The number of wells showing visible growth were noted, and OD_600_ was measured using a plate reader. Two controls were also set up in parallel. For the first control, primary competition was performed in the absence of trimethoprim such that there would be no enrichment of resistant-mutants. The second control had similar number of wild-type cells without the addition of mutant bacteria. This control would indicate the frequency of spontaneous resistant mutants emerging during the competition. OD_600_ values of 144 replicate competitions for each condition were plotted and compared.

### Sequence and phylogenetic analyses

To analyse the distribution of homologs of PhoQ and MgrB from *E. coli* across different bacterial species, the Pfam database [43] was initially mined. Since both proteins were restricted to the Order Enterobacterales further analyses were restricted to this group of bacteria. The phylogenetic relationships of different families under Order Enterobacterales was taken from Adelou et al., 2016 [44]. Type strains from each family were identified using the LPSN (List of Prokaryotic names with Standing in Nomenclature; https://lpsn.dsmz.de/) database [79, 80]. For family Enterobacteriaceae, a maximum likelihood phylogenetic tree was constructed using 61 bacterial species in MEGA-X [81], by using 16s rRNA gene sequences of the type strain of each genus as listed in LPSN [79, 80]. The presence or absence of PhoQ and MgrB in bacterial strains under consideration was determined by three methods: first, directly analysing genome annotation data available on NCBI, second, by performing BLASTn against the genome of the query organism (nucleotide similarity with wild-type *E. coli mgrB*-NCBI Gene ID: 946351 and *phoQ*-NCBI Gene ID: 946326), third, by performing NCBI BLASTp against the proteome of the query organism (amino acid similarity with wild-type *E. coli* MgrB-Uniprot: P64512 and PhoQ-Uniprot: P23837). A positive hit in at least one of the above searches was taken to mean that PhoQ/MgrB were present in the query organism.

### Gene expression analysis

#### Transcriptome analyses using RNA-sequencing

For RNA-seq, 1% of a saturated culture of wild type and trimethoprim resistant isolates Tmp^R^-A, Tmp^R^-B and Tmp^R^-C was inoculated into 3 mL LB in duplicate and incubated at 37 ºC for 3 hours with shaking at 180 rpm. Bacteria were pelleted down using centrifugation and resuspended in 1 mL of RNAlater (Invitrogen) for storage. Subsequent RNA extraction, sequencing and preliminary data analyses were performed by Redcliff Life Sciences (India). Total RNA was extracted using RNAeasy spin columns and quantitated using Qubit and Bioanalyzer. Bacterial rRNA was depleted using Ribozero kit. Bacterial mRNA was then reverse transcribed, library was prepared and quality control was performed using Tapestation platform. Paired end sequencing was performed on Illumina platform with read lengths of 150 bp read length. Processed reads were aligned to the reference genome (NZ_CP025268) using HISTAT2(version 2.1.0). Abundance estimation was done using featureCounts(version1.34.0). Differential gene expression (DGE) analysis was done by comparing the expression of individual genes trimethoprim-resistant strains to wildtype using DESeq2. The output of the DGE analyses were log_2_(fold change) values for each gene and P-values to test for statistical significance. Differentially expressed genes were classified into PhoP-regulated, RpoS-regulated and RpoD-regulated based on information available in RegulonDB [47, 48] and Ecocyc [46]. Sigma factors regulating differentially expressed genes were also obtained from RegulonDB [47, 48] and Ecocyc [46].

#### Gene specific quantitative RT-PCR

Quantitative RT-PCR was used to validate the findings of RNA-seq. For RNA extraction, appropriate bacterial strains were grown for 3 hours at 37 ºC and pelleted by centrifugation. Total RNA was extracted using TRIzol Reagent (Invitrogen, USA) and quantified spectrophotometrically. RNA quality was evaluated by electrophoresis on a 1% agarose gel and staining with ethidium bromide. Extracted RNA (20µg) was treated with recombinant DNase I (RNAse free) (Takara, Japan) and then cleaned-up using RNeasy spin column (Qiagen, Japan). Prepared RNA was stored at -80 ºC until further use. Reverse transcription reaction was set up using the cleaned-up RNA (~1.6µg) using PrimeScript™ RT reagent Kit (Takara, Japan). The RT reaction was carried out with buffering temperature at 25 ºC for 2 mins, cDNA synthesis at 37 ºC for 30 mins. The RT enzyme was inactivated at 85 ºC for 90 seconds. Prepared cDNA was serially diluted (10, 100 and 1000-fold) and semi-quantitative PCR were set-up using gene-specific primers (Table 2) to ensure the absence of contaminating genomic DNA contamination and checking for priming efficiency. cDNA was stored at -20 ºC until further use.

**Table 2.**
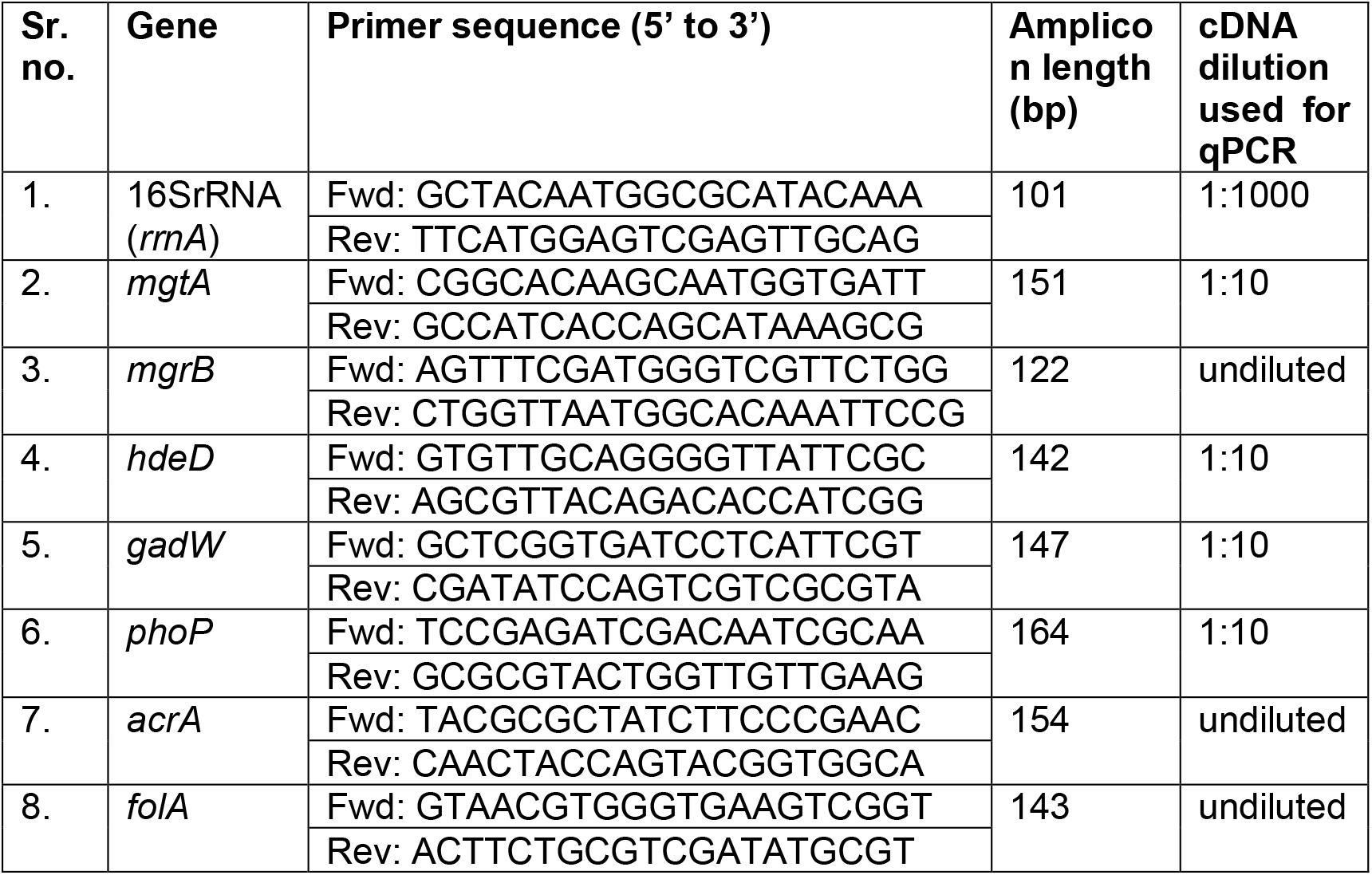
List of primer, cDNA dilution and expected product sizes for qPCR

Quantitative RT-PCR (qPCR) for target genes was performed using TB Green®Premix Ex Taq™ II (Tli RNaseH Plus) (Takara, Japan) and using gene-specific primers listed in Table 2. For each PCR, 20 µL reaction contained 10 µL of 2X TB Green Premix, 0.5 µl each of forward and reverse primers (10 µM stock), 8µl of nuclease-free water and 1 µL of appropriately diluted cDNA. The qPCR reactions were performed on an Eppendorf Realplex2 Mastercycler (Eppendorf, Germany) using a two-step protocol with an initial denaturation at 95 ºC for 30 seconds, 40 cycles of denaturation at 95 ºC for 5 seconds and annealing and extension for 30 seconds at 60 ºC. A melt curve analysis was performed after 40 cycles of qPCR. 16S rRNA was used as a normalizing internal control to calculate change in expression for all other genes. A known concentration of genomic DNA and its dilutions were used to construct a standard graph for each gene. Fold changes in gene expression for each gene were calculated with respect to the wild type using the standard graph method.

## Acknowledgements

Funding for this work was provided by the DBT/Wellcome Trust India Alliance Intermediate Fellowship awarded to NM. RV and MB are supported by fellowships from the Indian Institute of Science Education and Research, Pune, India. CY is supported by fellowship from the Department of Biotechnology, Government of India.

## Material and Data Availability

All materials such as bacterial strains generated in this study will be shared upon request. Sequencing datasets will be uploaded to Genbank soon (Project:PRJNA741586) and shared on demand if needed.

